# Methylation of *E. coli* Class I Release Factors Increases the Accuracy of Translation Termination *in vitro*

**DOI:** 10.1101/239822

**Authors:** Gürkan Korkmaz

## Abstract

Ribosomal protein synthesis (translation) is a highly accurate process. Translation termination, in particular, must be accurate to prevent truncated proteins. How this accuracy is achieved is not fully understood in all its details. Using an *E. coli in vitro* system, I explore novel mechanisms that contribute to the high accuracy of translation termination. By comparing the Michaelis-Menten parameters of methylated and non-methylated release factors on cognate and non-cognate codons. Post-translational methylation of a strictly conserved GGQ motif in class I release factors increases the accuracy of termination by up to 5-fold. This happens by increasing both the maximum rate of peptide release (*k_cat_*) and Michaelis-Menten constant (*K_M_*). Further, I demonstrate here that a non-methylated release factor acts like an uncompetitive inhibitor of enzyme reactions. Overall, this study shows that the methylation of class I release factors is a novel mechanism contributing to highly accurate translation termination.

**Abbreviations:**

RF: release factor
RC: release complex

## INTRODUCTION

Translation termination is an essential component of protein production. mRNA translation is terminated when a stop codon is translocated into the ribosomal A site. This new configuration is now called a release complex (RC). The subsequent binding of the class I release factor induces peptide release. In nearly all life forms, the codons UAA, UAG, and UGA are used as termination signals, with some exceptions (Ivanova et al., 2014). In bacteria, the class I release factors are RF1 and RF2. Both class I release factors possess a strictly conserved glycine-glycine-glutamine (GGQ) motif, which is essential for the ester bond hydrolysis by which the peptide is released from the tRNA.

Stop codon recognition is highly accurate *in vitro* (Freistroffer et al., 2000) and *in vivo* (Jørgensen et al., 1993), but our understanding of how this accuracy is achieved is still rudimentary. The accuracy of a reaction is defined as the ratio of the efficiency of a cognate to a non-cognate reaction (Freistroffer et al., 2000). The efficiency is determined by the ratio of the maximum rate of the reaction (*k_cat_*) to the concentration at half *k_cat_* (*K_M_*). When the ratio of cognate and non-cognate reaction is one, accuracy is non-existent, meaning no discrimination between the cognate and non-cognate substrate are observable.

Methylation of the glutamine in the GGQ motif has been shown to increase *k_cat_* (Indrisiunaite et al., 2015). Methylation happens via the hemK protein, which modifies glutamine 253 (*E. coli* numbering) to N^5^-methyl-glutamine (Heurgue-Hamard et al., 2002). However, its effect on *K_M_* is unknown.

Highly conserved recognition motifs of RF1 and RF2 are located in the vicinity of the stop codons, but about 75Å distant from the catalytic center (Korostelev, 2011). These motifs (PxT in RF1 and SPF in RF2) were thought to be responsible for the specificity of stop codon recognition by the release factors and therefore named ‘tripeptide anticodons’, analogous to the anticodons of tRNAs (Ito et al., 1996, 2000).

Class II release factor RF3 was shown to affect accuracy partly (Freistroffer et al., 2000). Further, it was shown that depletion of the ribosomal protein L11 from an RC increases the accuracy of translation termination with RF1 but not RF2 (Bouakaz et al., 2006). The accuracy of translation termination is mostly thought to be due to the extensive interaction network between class I release factor, stop codon, and rRNA (Sund et al., 2010).

The strictly conserved GGQ motif is essential for ester bond hydrolysis, as shown by studies where mutations within the GGQ motif disrupted translation termination *in vitro* (Zavialov et al., 2002) and *in vivo* (Frolova et al., 1999; Seit-Nebi et al., 2001). However, in *E. coli* different mechanisms seem to be at play for several reasons. The conserved methylation of the GGQ motif glutamine to N^5^-methyl-glutamine is not essential in *E. coli*, as the responsible enzyme for methylation RF1/RF2, hemK, is not essential for cell viability (Heurgue-Hamard et al., 2002; Pannekoek et al., 2005). Also, release factors where the GGQ motif is altered to GGA are still able to induce translation termination *in vitro* (Shaw and Green, 2007; Zavialov et al., 2002). On the other hand, methylation of the class I release factors is essential for cell viability in minimal media (Mora et al., 2007). As mentioned earlier, methylation increases the maximum rate of peptide release (feat) (Indrisiunaite et al., 2015), but the magnitude of the *k_cat_* increase was shown to depend on the nature of the amino acid on the tRNA occupying the P site (Pierson et al., 2016).

The actual ester bond hydrolysis occurs via a coordination of a nucleophilic hydroxide molecule by the GGQ motif into the PTC. Crystal structures of class I release factors indicate a difference between the positioning of the methylated and non-methylated glutamine residue within the PTC (Pierson et al., 2016).

Although RF1 and RF2 share high sequence and structural similarity, they have different codon specificities. RF1 recognizes the stop codons UAA and UAG, whereas RF2 recognizes the stop codons UAA and UGA (Scolnick et al., 1968).

## RESULTS & DISCUSSION

In this study, I investigated the methylation of the GGQ glutamine as an additional factor that might enhance the accuracy of translation termination *in vitro*. The accuracy of a reaction is defined as the ratio of the efficiency (*k_cat_* / *K_M_*) of a cognate to a non-cognate reaction (Freistroffer et al., 2000). When the ratio of cognate and non-cognate reaction is one, accuracy is non-existent, meaning no discrimination between the cognate and non-cognate substrate are observable. I systematically determined the *k_cat_* and *K_M_* parameters for release factors on various purified release complexes (Korkmaz and Sanyal, 2017) containing cognate and non-cognate codons in A site and compared the resulting accuracy with the reactions involving methylated (mRF1 or mRF2) and non-methylated (RF1 or RF2) release factors (Table 1 and Figure 1 and 2). UAA was used as the cognate codon in all experiments, while UAG and UGA were used as the non-cognate substrates for RF1 and RF2 respectively since these codons allow discrimination similar to non-cognate sense codons (Freistroffer et al., 2000). The determined cognate *k_cat_* parameters were in agreement with previous data for non-methylated release factors (Bouakaz et al., 2006; Freistroffer et al., 2000) and also with recent studies that used methylated release factors (Indrisiunaite et al., 2015; Pierson et al., 2016). I observed a 4.5-fold increase in the maximum rate of peptide release for mRF1 on its cognate UAA codon and a 5-fold increase in the maximum rate of tritium-labeled fMet (^3H^fMet) release on the non-cognate codon UGA. However, it should be noted that for the cognate reactions, the *K_M_* values for reactions with either mRF1 or RF1 were below the detection limit of 5 nM. Thus, the real difference in *K_M_* might be much larger than what could be measured in these experiments. In Table 1 the estimated *K_M_* parameters are listed. The *K_M_* value form RF1 on UGA decreased by 3-fold, from 164 nM to 48 nM, when methylation was missing. These resulted in an accuracy of 25 for RF1 on UGA. When RF1 was methylated, the accuracy increased to 90 (an increase of over 3.5-fold) (Table 1).

**Table 1.**
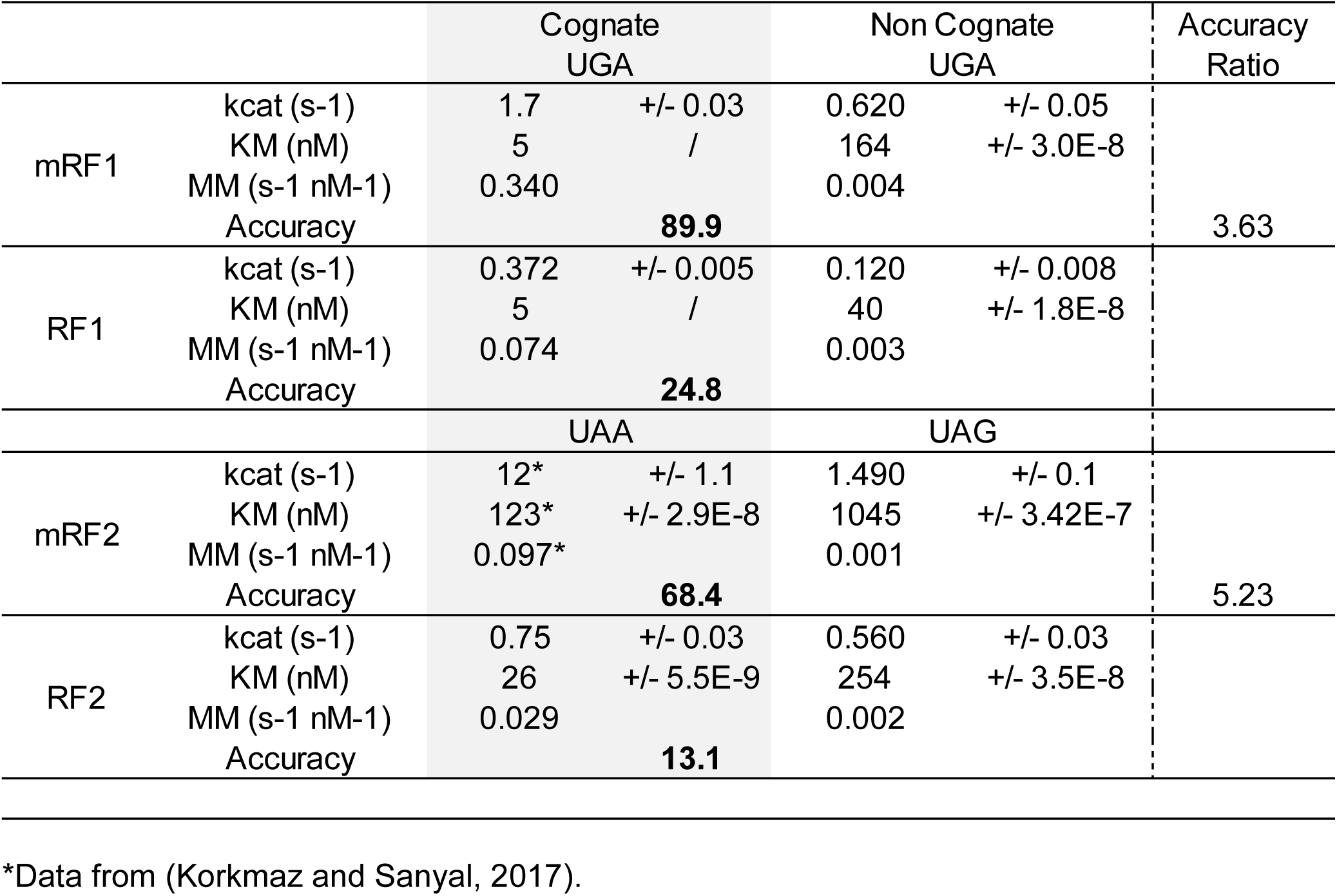
Comparison of Michaelis-Menten parameters and accuracy measurements of methylated and non-methylated class I release factors. The *E. coli* K12 genes encoding RF1 (*prfA*) and RF2 (*prfB*) were already available as clones in the pTRC and pET11a vectors, respectively. The final construct pET11a-prfB encoded a prfB^T246A^ variant to enable the overexpression of an otherwise toxic RF2 (Uno et al., 1996). Both constructs contained C-terminal His6-tags. The release factor methyltransferase *hemK* (gene: *prmC*), was available as a clone in pACDuet1; the protein contained no tag. All constructs were under the control of an Isopropyl p-D-1-thiogalactopyranoside (IPTG) inducible *lac* promoter. Plasmids pTRC-prfA and pET11a-prfB were introduced separately into *E. coli* BL21 (DE3)Gold cells (Agilent Technologies), already harbouring pACDuet1-prmC, via chemical transformation. Protein production was induced with 1 mM IPTG in exponential phase cultures grown at 37 °C in LB medium. To allow full methylation of the class I release factors, cells were further grown for 4 h. The bacteria were lysed using a French press and purified using HisTrap nickel affinity chromatography (GE Healthcare) per manufacturer’s instructions. Purified proteins were concentrated, dialyzed against 1 L of Polymix Buffer (pH 7.5) (Indrisiunaite et al., 2015; Jelenc and Kurland, 1979), with one buffer exchange after 3 h, and stored at - 80 °C until use. Methylation of the release factors was verified by mass spectrometry as described earlier (Heurgue-Hamard et al., 2002). For the *in vitro* assays, the experiments were prepared and performed as described earlier (Korkmaz and Sanyal, 2017) with the following mRNAs 5’ GGG AAU UCG GGC CCU UGU UAA CAA UUA AGG AGG UAU ACU <u>AUG</u> STOP CUG CAG (A)_21_ 3’ (the start codon is underlined, and STOP indicates the position of the stop codon).

**Figure 1.**
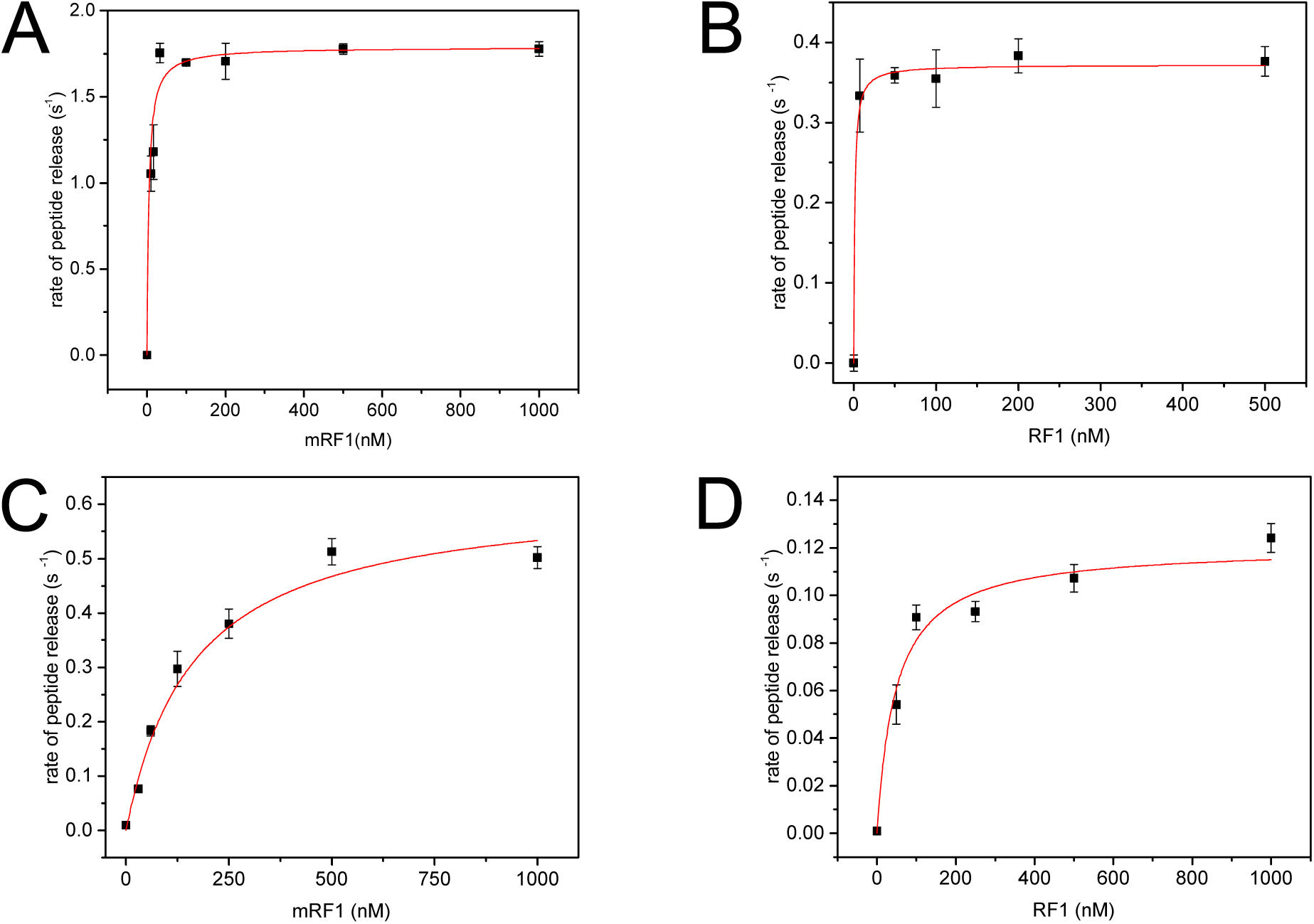
Michaelis-Menten graphs of class I release factors on cognate and non-cognate substrates. (A) mRF1 on UAA (B) RF1 on UAA (C) mRF1 on UGA (D) RF1 on UGA

**Figure 2.**
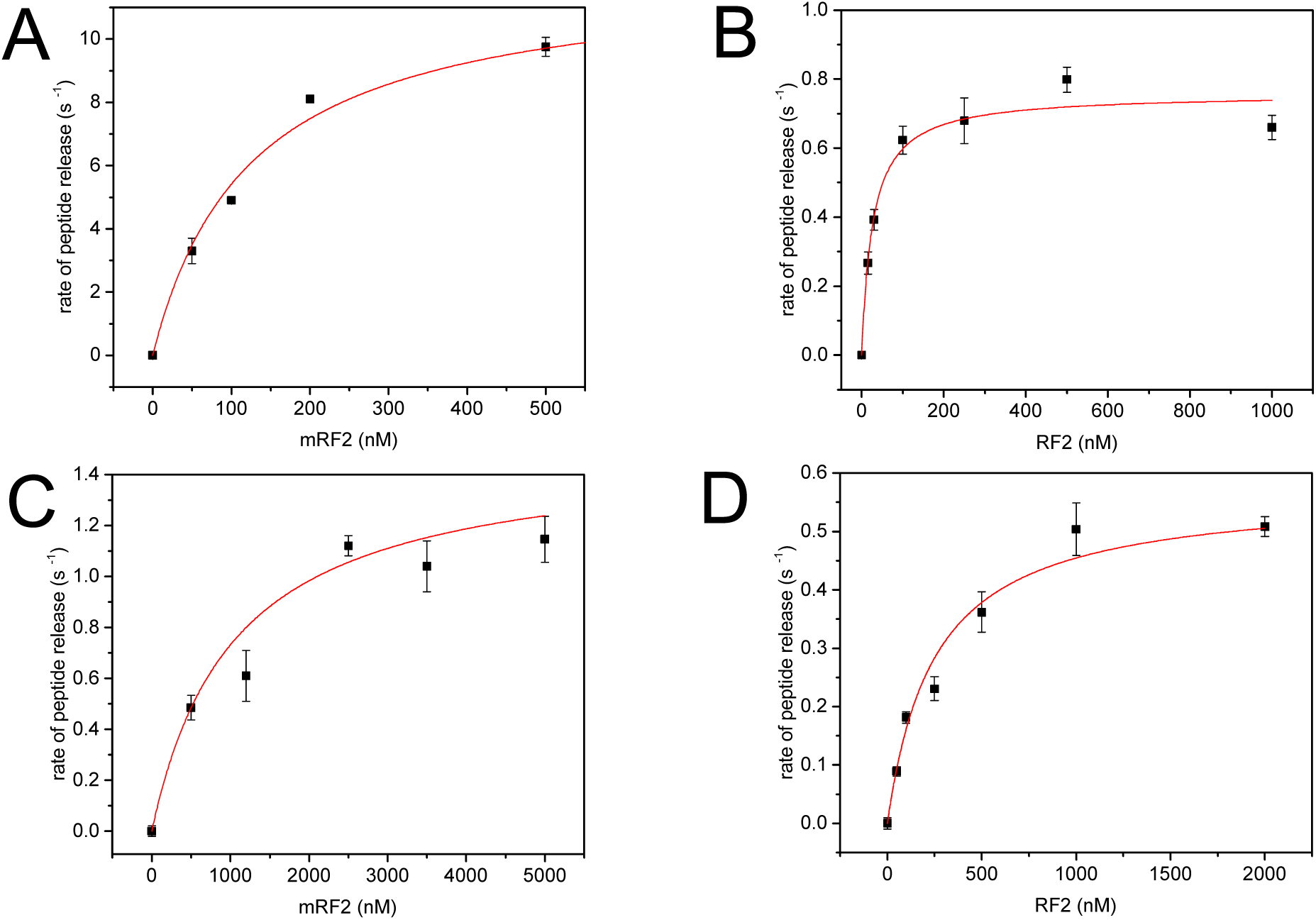
Michaelis-Menten graphs of class I release factors on cognate and non-cognate substrates. (A) mRF2 (Korkmaz and Sanyal, 2017) on UAA (B) RF2 on UAA (C) mRF2 on UAG (D) RF2 on UAG

**Figure 3.**
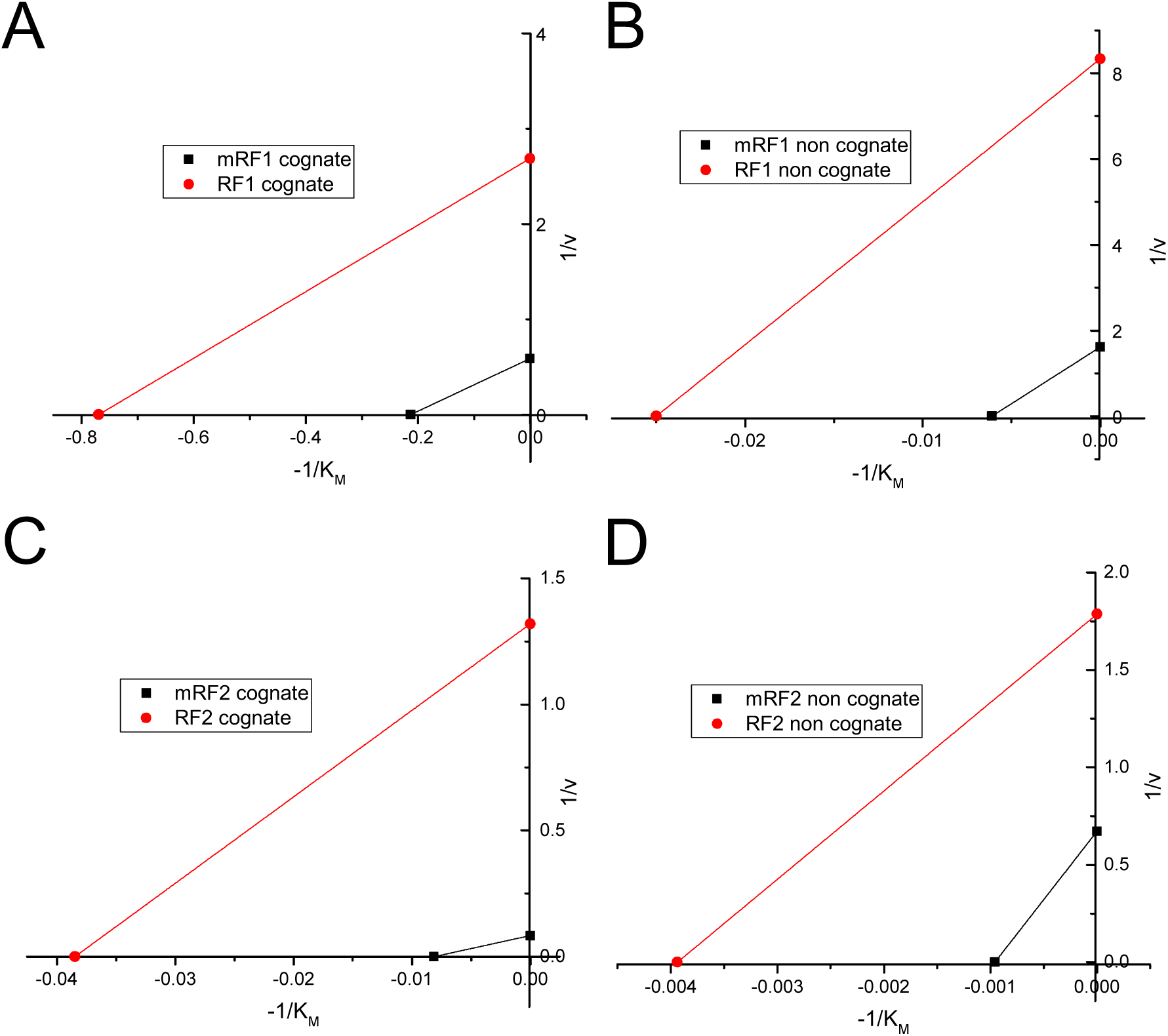
Lineweaver-Burk plots of kinetic data from Table 1. (A) Lineweaver-Burk plot comparison of methylated and non-methylated RF1 kinetic data on UAA. For illustration purposes, the *K_M_* value for RF1 was evaluated as observed in Table 1. (B) Comparison of mRF1 and RF1 on UAG. (C) Comparison of mRF2 and RF2 on UAA. (D) Comparison of mRF2 and RF2 on UAG. All *K_M_* values are given in nM.

The effect of methylation on the translation termination accuracy was even more pronounced with RF2 on the non-cognate UAG codon. With methylation, the *k_cat_* increased 2.6-fold for the non-cognate substrate and nearly 16-fold for the cognate substrate (UAA). The accuracy of translation termination increased 5-fold, from 13 to 68, when RF2 was methylated.

As mentioned earlier, it is not surprising that the maximum rate of ^3H^fMet release increased in all cases; however, unexpectedly, the *K_M_* parameters were also affected by release factor methylation. For RF1 and mRF1, the change in *K_M_* (Table 1) could not be unambiguously determined in the experimental setup as both values were below the detection limit, but an approximately 3-fold difference in *K_M_* was observed for the non-cognate reading of UGA; methylation of RF1 resulted in an increase of the *K_M_* value. More pronounced changes in *K_M_* were observed for RF2 and mRF2. For cognate stop codon recognition, a nearly 5-fold increase in *K_M_* was observed when RF2 was methylated (26 nM for RF2 and 123 nM for mRF2); a 4-fold increase was seen in the case of the non-cognate substrate UAG (251 nM for RF2 and 1045 nM for mRF2).

When hemK is knocked out, cells have compromised fitness, yet they are viable (Mora et al., 2007). The knockout of hemK consequently leads to the lack of methylation in class I RFs and as shown here. The protein hemK becomes essential when cells are grown in minimal media (Mora et al., 2007). In the light of the data presented here, one could explain this phenotype by the fact of the lower accuracy of RFs than the slowed down peptide release. In minimal media, ribosomes are likely to pause on codons due to the amino acid limitation (Li et al., 2012). Inaccurate RFs could bind to these complexes and induce premature termination.

Interestingly, the Lineweaver-Burk plots of the methylated and non-methylated release factor kinetics data revealed that the non-methylated release factors behave similarly to uncompetitive inhibitors in enzyme reactions (Figure 3). However, the mechanism of the inhibition or details of any mechanistic similarity to uncompetitive enzyme inhibition remains obscure.

Overall, I herein presented evidence that methylation of class I release factors contribute to the accuracy of translation termination. This is surprising by the fact that a post-translational modification, far from the initial binding site, effects the binding constant *K_M_*.

Further studies, unraveling the potential *in vivo* aspects of this finding are imminent, especially in light of the ribosome rescue phenomenon. This study demonstrates that release factor methylation plays an additional biological role beyond the known increase of the maximum rate of peptide release. This previously unknown variable can explain how the outstanding accuracy of translation termination is achieved in the absence of any proofreading steps.

## ACKNOWLEDGEMENTS

GK was funded partly by a scholarship from the Sven and Lilly Lawski Foundation for Scientific Research. GK would like to acknowledge Marina Fridman from the Champalimaud Centre for the Unknown, Lisbon, Portugal, for proofreading an early version of the manuscript. Further Suparna Sanyal from the Department of Cell and Molecular Biology, Uppsala University, Sweden, is acknowledged for grant acquisition that supported the experimental part of this work (Swedish Research Council 2013-8778, 2014-4423, 2008-6593 [Linnaeus grant to Uppsala RNA Research Center], and the Knut and Alice Wallenberg Foundation to RiboCORE [KAW2011.0081]).

